# Epithelial Cell-like Elasticity Modulates E-cadherin Adhesion Organization

**DOI:** 10.1101/2020.05.22.111443

**Authors:** Mohamad Eftekharjoo, Siddharth Chatterji, Venkat Maruthamuthu

**Affiliations:** Department of Mechanical & Aerospace Engineering, Old Dominion University, Norfolk, VA

## Abstract

E-cadherin adhesions are essential for cell-to-cell cohesion and mechanical coupling between epithelial cells and reside in a micro-environment that comprises the adjoining epithelial cells. While E-cadherin has been shown to be a mechanosensor, it is unknown if E-cadherin adhesions can differentially sense stiffness within the range of that of epithelial cells. A survey of literature shows that epithelial cells’ Young’s moduli of elasticity lie predominantly in the sub kPa to few kPa range, with cancer cells often being softer than non-cancerous ones. Here, we devised oriented E-cadherin-coated soft silicone substrates with sub kPa or few kPa elasticity, but with similar viscous moduli, and found that E-cadherin adhesions differentially organize depending on the magnitude of epithelial cell-like elasticity. Linearly shaped E-cadherin adhesions associated with radially oriented actin, but not irregularly shaped E-cadherin adhesions associated with circumferential actin foci, were much more numerous on 2.4 kPa E-cadherin substrates compared to 0.3 kPa E-cadherin substrates. However, the total amount of E-cadherin in both types of adhesions taken together was similar on the 0.3 kPa and 2.4 kPa E-cadherin substrates, across many cells. Furthermore, while the average extent of nuclear recruitment of the mechanoresponsive transcription factor YAP on the 0.3 kPa E-cadherin substrate was undiminished relative to that on the 2.4 kPa substrate, a fraction of cells on the softer substrate displayed relatively high levels of YAP nuclear recruitment. Our results show that E-cadherin adhesions can be regulated by epithelial cell-like elasticity and have significant implications for disease states like carcinomas characterized by altered epithelial cell elasticity.

## Introduction

E-cadherin is a cardinal epithelial cell-cell adhesion molecule that is essential for proper morphogenesis as well as for the maintenance of the architecture of adult epithelial tissues [1, 2]. At the micro-scale, E-cadherin adhesions are collectives of many trans-interacting E-cadherin cis-dimers/clusters from the surface of neighboring epithelial cells [3]. They form functional units that intimately couple to the underlying actin cytoskeleton to integrate the contractile and adhesive responses of epithelial cells [4–6]. Understanding both the biochemical and biophysical aspects of E-cadherin adhesions is essential to deciphering their role in larger cell collectives and tissues. Importantly, E-cadherin adhesions have been shown to be bonafide mechanosensors [7], much like integrin-based adhesions [8]. While integrin-based adhesions are known to be sensitive to elasticity within the range of that of the extra-cellular matrix (ECM) [8], it is unclear if E-cadherin-based adhesions can differentially sense elasticity within the range of that of epithelial cells (that enclose them). Data from multiple human epithelial cells obtained using atomic force microscopy have shown that epithelial cell elasticities (Young’s moduli) hover around 1 kPa [9].

E-cadherin adhesions are, naturally, primarily studied using epithelial islands or monolayers that have extensive cell-cell contacts. However, E-cadherin adhesions at epithelial cell-cell contacts exist among a complex mileu of other cell-cell adhesion systems. Epithelial cell-cell contacts consist of several types of cell-cell junctions including tight junctions, adherens junctions, desmosomes and gap junctions [10]. Adherens junctions themselves consist of not only E-cadherin adhesions, but also other types of cell-cell adhesions such as those mediated by nectins. Several of these adhesion molecules are regulated by the common framework of the cell’s cytoskeleton, including F-actin. To enable the specific probing of cadherin, several groups have utilized cadherin-coated substrates [11–14]. Glass surfaces coated with the extra-cellular region of E-cadherin fused to the Fc region (E-cadherin-Fc) have especially been used to study biochemical events initiated specifically by E-cadherin adhesion [15]. Such flat E-cadherin surfaces also enable easier imaging of adhesions compared to native epithelial cell-cell contacts that have a more complex topology. Flexible E-cadherin-coated substrates have specifically been used to understand E-cadherin mechanobiology [16, 17]. Previous reports have shown that E-cadherin adhesions can distinguish between elastic moduli within the tens of kPa range [16] (similar to N-cadherin [18]) as well as between kPa and MPa elastic moduli [17]. However, it is unclear if E-cadherin adhesions can differentially sense elasticity within the range of that of epithelial cells, i.e., in the sub-kPa to few kPa range [9]. To our knowledge, prior studies that used E-cadherin-coated surfaces have not reported on the formation of E-cadherin adhesions corresponding to this crucial range of elasticity.

E-cadherin is a well-known tumor suppressor but is continued to be expressed in many types of cancers [19]. It has been shown that modulation of E-cadherin adhesion may play a key role in cancer progression [20]. In this context, it has been proposed [20] that inside-out signaling can modulate E-cadherin adhesions. Since human epithelial cancer cells are often softer than normal cells [9], it is also possible that E-cadherin mechanosensing of cell elasticity may be at play. We noticed that many, though not all, studies that compared normal or benign human epithelial cells with cancerous ones from the same tissue of origin reported a similar trend: the Young’s modulus was 2 kPa for normal vs 0.5 kPa for cancer cells from lungs [21], 2 kPa for normal vs 0.5 kPa for cancer cells from breasts [21], 2.5 kPa for normal vs 0.5-1.1 kPa for cancer cells from ovaries [22], 2.2 kPa for normal vs 1.4 for cancer cells from thyroid [23] and 2.8 kPa for normal vs 0.3-1.4 kPa for cancer cells from prostate [24]. Due to the preponderance of human epithelial cell elastic moduli of either sub kPa or few kPa magnitude, we sought to test if E-cadherin adhesions may differentially sense/respond to epithelial-cell like elasticities in this range. We first developed a biomimetic E-cadherin soft substrate by modifying prior approaches and then proceeded to employ it in answering the question of E-cadherin sensing of epithelial cell-like stiffness as well as possible downstream consequences.

## Materials and Methods

### Cell culture

Human colon epithelial cells (C2BBe, a sub-clone of caco-2) were cultured overnight in Dulbecco’s modified Eagle’s medium (Corning Inc., Corning, NY) supplemented with L-glutamine, sodium pyruvate, 1% penicillin/streptomycin, and 10% fetal bovine serum (Corning Inc., Corning, NY) under 5% CO2 at 37 °C. Before each experiment, cells were detached from cell culture dishes using a trypsin-free chelator-based cell dissociation reagent (Versene, Thermo Fisher Scientific, Waltham, MA). The cells were seeded on samples in the same media as above, but without serum, for 2 hours under 5% CO2 at 37 °C.

### Biomimetic E-cadherin soft substrate preparation

Soft silicone (GEL8100, NuSil Silicone Technologies, Carpinteria, CA, USA) was prepared by mixing the base and crosslinker components (labelled A and B) in the ratio 2:3 or 2:7 by weight. The storage and loss shear moduli of each formulation was characterized using an HR-2 Discovery rheometer (TA Instruments, New Castle, DE) in a parallel plate geometry. To prepare the substrates, about 100 μL of soft silicone was first pipetted onto a 22 mm x 22 mm glass coverslip and cured for 1 hour at 100 °C on a hot plate. The substrate was then exposed to 305 nm UV light (UVP crosslinker, Analytik Jena AG, Upland, CA) for 5 minutes. Protein A (Prospec, Rehovot, Israel) was then coupled to the substrate by incubation with a mixture of 0.2 mg/mL protein A, 10 mg/mL EDC (1-ethyl-3-(3-dimethylaminopropyl) carbodiimide hydrochloride) and 5 mg/mL sulfo-NHS (N-hydroxysulfosuccinimide) for 1 hr at room temperature. After washing with PBS (with calcium), the substrate was incubated with 0.1 mg/ml recombinant E-cadherin-Fc for 2 hours (Sino Biological, Beijing, China). Afterwards, the sample was washed with PBS (with calcium) and incubated with 1 mg/ml Fc fragment (Jackson ImmunoResearch, West Grove, PA) for 1 hour. The sample was again washed with PBS (with calcium) before cell plating.

### Immunofluorescence and imaging

Cells were permeabilized and fixed in buffer C (10 mM MES (2-morpholinoethanesulfonic acid), 3 mM MgCl2 and 138 mM KCl; pH 6.8) with 4% paraformaldehyde, 1.5% (w/v) bovine serum albumin and 0.5% (v/v) Triton-X for 15 min. The cells were incubated overnight at 4 °C with primary antibodies and for 1 hour at room temperature with secondary antibodies. Primary antibodies used were anti-β-catenin (Clone 14, 610154, BD Biosciences, San Jose, CA), anti-paxillin (Y113, ab32084, Abcam, Cambridge, UK), anti-E-cadherin (DECMA-1, sc-59778, Santa Cruz Biotechnology, Dallas, TX) and anti-YAP (sc-101199, Santa Cruz Biotechnology, Dallas, TX). Alexa Fluor 488 conjugated phalloidin was from Thermo Fisher Scientific, Waltham, MA and DAPI was from Biotium, Hayward, CA. Secondary antibodies were from Jackson ImmunoResearch, West Grove, PA. All images were taken using a Leica DMi8 epifluorescence microscope (Leica Microsystems, Buffalo Grove, IL) with 10x, 20x or 40x objectives and a Clara cooled CCD camera (Andor Technology, Belfast, Northern Ireland).

### Image analysis

To compare E-cadherin coating densities on the silicone surfaces, we used ImageJ [25] to extract the mean intensity values of the regions with deposited E-cadherin (as immunostained with anti-E-cadherin antibody). For E-cadherin adhesion analysis, discrete E-cadherin adhesions ~0.5 μm^2^ or larger were manually segmented using CellProfiler (Version 3.1.9) [26], using a wacom intuos pen tablet. Individual adhesion intensity and shape features were also extracted using CellProfiler. The eccentricity of an adhesion was obtained by first fitting an ellipse with the same second moment as the adhesion and then computing the ratio of the distance between the foci of the ellipse and its major axis length. To quantify YAP nuclear recruitment, we segmented DAPI stained nuclei and actin stained cells and extracted the mean YAP intensity in the nuclear and cytoplasmic regions. The ratio of the above two mean YAP intensities yielded a measure of YAP nuclear recruitment for each cell [27].

### Statistical analysis

A two-tailed Student’s t-test was used with cell area data. A Welch’s t-test was used with the YAP nuclear recruitment data. MATLAB (MathWorks, Natick, MA) or Excel (Microsoft, Redmond, WA) were used to carry out the statistical analysis.

## Results and Discussion

The broad question we were interested in was whether E-cadherin adhesions can sense, and be affected by, an elastic microenvironment that mimics that of the epithelial cells that enclose them. E-cadherin adhesions reside on the surface of epithelial cells, in close proximity to the cell cortex. Thus, we reasoned that values of epithelial cell stiffness reported in literature using methods such as atomic force microscopy would be most relevant to E-cadherin mechanosensing – as opposed to methods which measure the elasticity deep in the cytoplasm that don’t involve contributions from the cell cortex [28]. A survey [9] of several studies that used atomic force microscopy to measure normal and cancer human epithelial cells of the same tissue origin shows (fig. 1A) that (i) human epithelial cell stiffness lies in the sub kPa to few kPa range and (ii) cancer cell stiffness is typically lower than that of normal cells. It has been previously shown that E-cadherin adhesions can sense applied forces [29, 30] as well as tens of kPa stiffness [16]. Considering the overall distribution of human epithelial cell stiffness (normal as well as cancerous) in fig. 1A, we asked whether E-cadherin adhesions may differentially sense elasticity within this range. To attain this elasticity range, we tested soft silicones of various compositions and came up with ones (fig. 1B,C) that had Young’s moduli (E ~ 3G’, where G’ is the storage modulus (elastic component)) of sub kPa (0.3 kPa) and few kPa (2.4 kPa) magnitude, while simultaneously having lower loss moduli (viscous component) of similar magnitude (tens of Pa) for either soft silicone (fig. 1B,C). Notably, these substrates have a loss modulus that is a fraction (~0.1-0.5) of the storage modulus, similar to that reported for epithelial cells previously [31].

**Figure 1.**
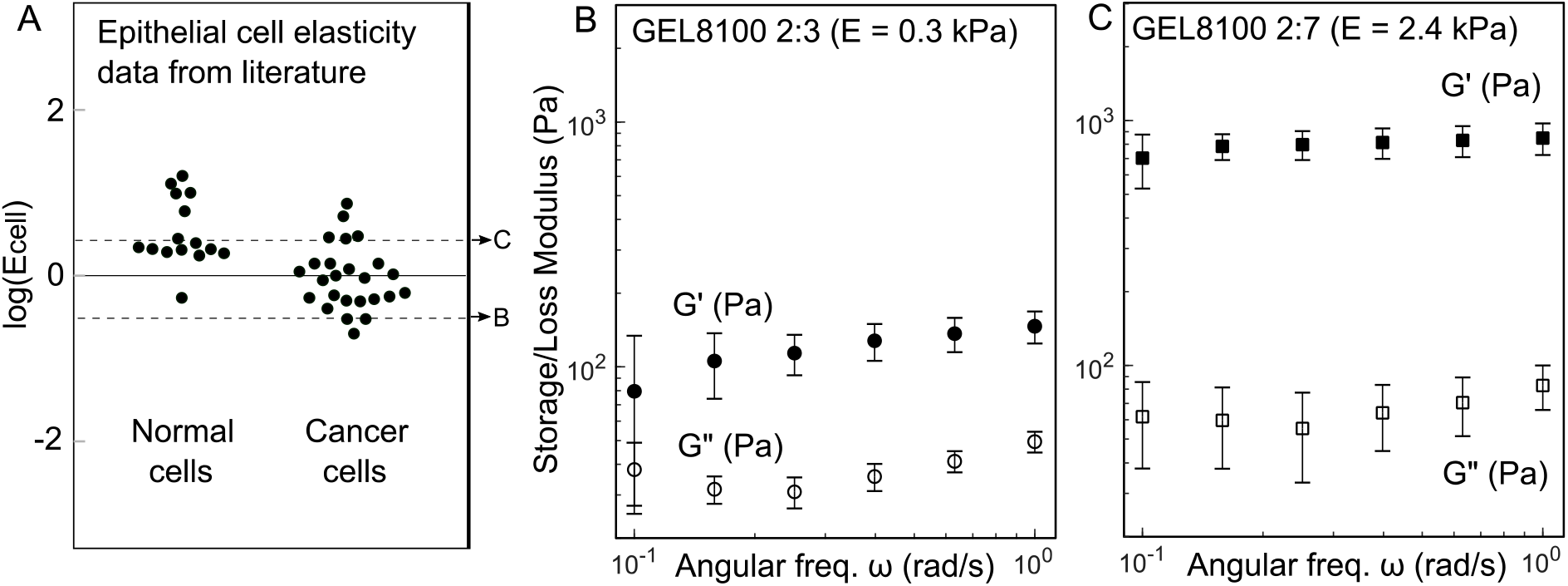
Soft silicone substrates with epithelial-cell like elasticity. (A) The Young’s moduli of various human normal/benign and cancer epithelial cells (Ecell) reported in literature, with Ecell in kPa. Note that Log10(Ecell) = 0 corresponds to Ecell of 1 kPa. Dotted lines correspond to the Young’s moduli of the soft silicones in (B) and (C). (B, C) Rheology data of 2:3 (B) and 2:7 (C) GEL8100 soft silicone corresponding to Young’s moduli E of 0.3 and 2.4 kPa, respectively. The storage (G’) and loss (G”) moduli of the soft silicones are shown as a function of angular frequency. Each data point is the mean ± s.d from 3 independent experiments. For incompressible soft silicone, E~3G’ with G’ being the average of G’ for ω = 0.1 to 1 rad/s.

To coat the soft silicone substrates with E-cadherin, we chose the strategy of oriented immobilization of Fc-tagged E-cadherin – based on earlier E-cadherin biophysical [32] and biochemical [33] studies (Fig. 2A). Here, the soft silicone was first coated with protein A (using EDC/sulfo-NHS chemistry) and then E-cadherin was immobilized onto protein A. Importantly, a similar strategy was shown recently to mimic lateral E-cadherin-based cell-cell junctions effectively [34]. Using immunofluorescence, we first checked that the density of E-cadherin on the 0.3 kPa and 2.4 kPa soft silicone substrates were similar (3360±340 and 3450±260 a.u, respectively; data from 2 independent samples for each substrate). We then plated human epithelial cells (C2BBe) on either substrate in the complete cell culture medium with serum. We found that the cells adhered, spread and formed E-cadherin adhesions (marked by β-catenin, since β-catenin binds to the cytoplasmic region of E-cadherin at a 1:1 stoichiometric ratio [35]) - but each cell also formed focal adhesions (marked by paxillin), as revealed by immunofluorescence (data not shown). To unambiguously attribute cell response to E-cadherin adhesion, we wanted to preclude the formation of focal adhesions. Use of integrin blocking antibodies used previously [36] did not prove successful for us. We reasoned that extra-cellular matrix components present in the serum could be contributing to the formation of (in this context, unwanted) focal adhesions. We then plated the cells in serum-free media and found that focal adhesion formation was considerably reduced, with a majority of cells that did not form any focal adhesions. Only cells with E-cadherin adhesions and no focal adhesions (as checked with each cell by immunofluorescence) were considered for all data and analyses that follow.

**Figure 2.**
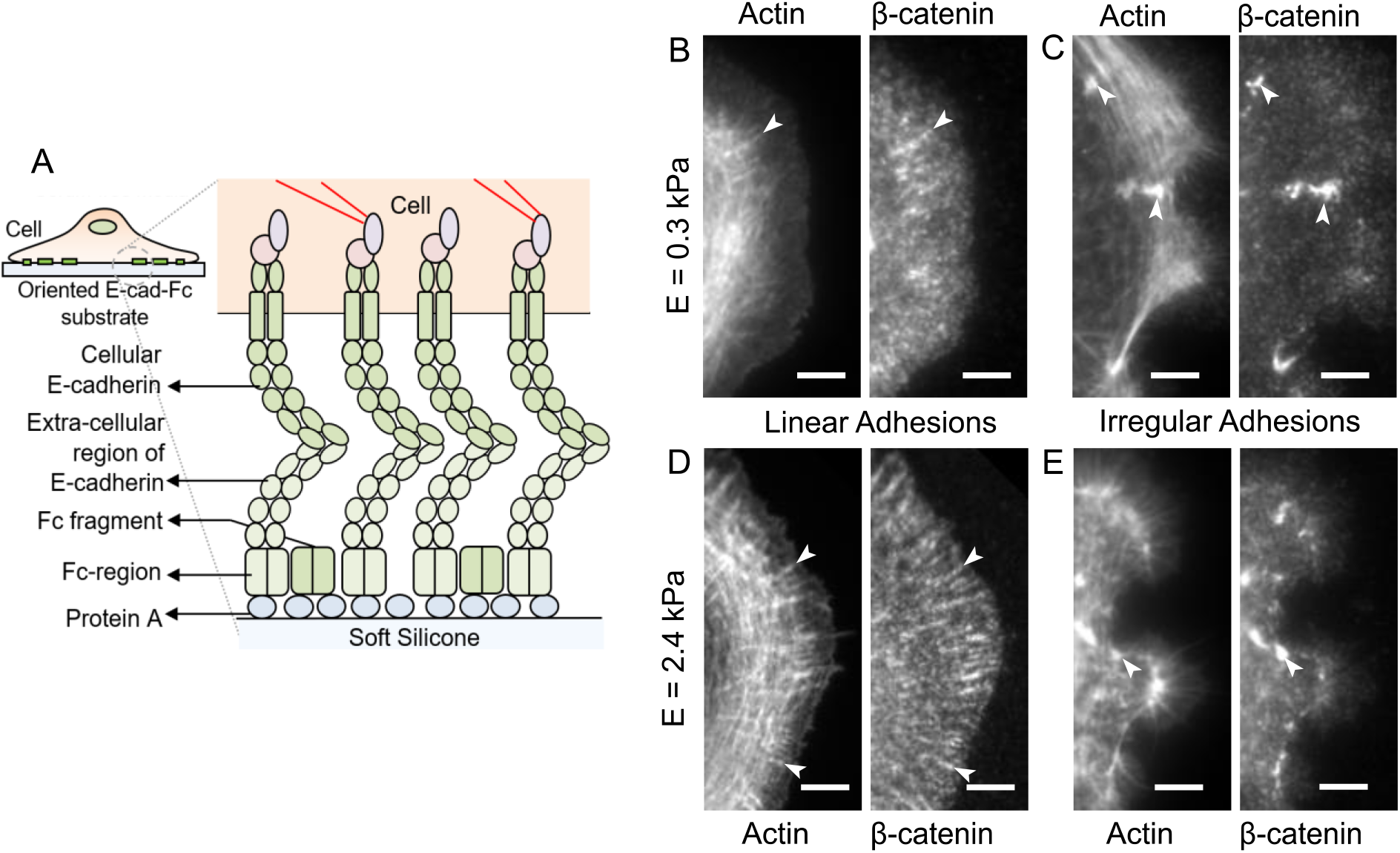
Actin-associated E-cadherin adhesions on sub kPa and few kPa E-cadherin substrates. (A) Schematic depiction of an epithelial cell adherent on an oriented, immobilized E-cadherin soft silicone substrate. (B-E) Immunofluorescence images of C2BBe epithelial cells adherent on 0.3 kPa (B, C) or 2.4 kPa (D, E) E-cadherin substrates, stained with phalloidin (to mark actin) and β-catenin (to mark E-cadherin adhesions). Linearly shaped, radial actin-associated E-cadherin adhesions (B, D) and irregularly shaped, circumferential actin foci-associated E-cadherin adhesions (C, E) are shown for both substrates. Arrow heads in the immunofluorescence images indicate representative radial actin or the associated linear E-cadherin adhesions (B, D) as well as representative circumferential actin foci and the associated irregular E-cadherin adhesions (C, E). All scale bars are 5 μm.

We first noted that the cell spread area on the 0.3 kPa and 2.4 kPa E-cadherin substrates were similar (1520±610 μm^2^ and 1470±450 μm^2^, respectively; p = 0.45, for 116 cells (0.3 kPa) and 145 cells (2.4 kPa) pooled from 8 independent experiments), unlike in studies where higher elasticity values were considered [16, 17]. The cells on both 0.3 kPa and 2.4 kPa E-cadherin substrates displayed a ‘background’ of diffraction-limited nascent E-cadherin adhesions throughout as well as larger discrete E-cadherin adhesions of >~0.5 μm^2^ (referred to simply as E-cadherin adhesions henceforth). Two kinds of E-cadherin adhesions were observed via immunofluorescence (Fig. 2B-E): linearly shaped adhesions associated with radially oriented Factin structures (Fig. 2B,D) and irregularly shaped adhesions associated with actin foci along circumferential F-actin structures (Fig. 2C,E). These two types of observed E-cadherin adhesions are henceforth referred to as linear and irregular adhesions. The radial actin bundles (reminiscent of dorsal stress fibers [37] for cells adherent on ECM), with which linear adhesions were associated, also appeared well-integrated with circumferentially oriented actin bundles (fig. 2B,D) (the latter reminiscent of transverse arcs [37] for cells adherent on ECM). The actin foci along circumferentially oriented actin structures, with which irregular adhesions were associated, were micron-scale regions of high F-actin intensity (fig. 2C,E) and such foci are not typically observed for cell adhesion to the ECM. The circumferential actin bundles (with which the radial actin appeared integrated), as well as the actin foci, were phosphomyosin positive, as revealed by immunofluorescence (data not shown). Both linear and irregular adhesions were largely spatially segregated from each other and were present in different cells to different extents based on the prevalence of radial actin or actin foci along circumferentially oriented actin structures.

We segmented the E-cadherin adhesions using CellProfiler [26] and determined the number, shape and intensity of E-cadherin adhesions of either kind on the 0.3 kPa and 2.4 kPa substrates. The eccentricities (shape factors) of the linear adhesions on the 0.3 and 2.4 kPa substrates were 0.92±0.04 and 0.91±0.06, respectively (insets in fig.3A,C). Note that the eccentricity of a line is 1 whereas that of a circle is 0, showing that the linear adhesions were indeed quite linearly shaped. In contrast, the eccentricities of the irregular adhesions on 0.3 and 2.4 kPa substrates were lower and more broadly distributed (0.78±0.16 and 0.79±0.15, respectively - insets in fig.3B,D). Overall, over twice as many E-cadherin adhesions formed on the 2.4 kPa substrate when compared to the 0.3 kPa substrate, in a similar number of cells (58 cells on the 0.3 kPa substrate and 59 cells on the 2.4 kPa substrate, pooled from 3 independent samples each). When resolved into linear and irregular adhesions, we found that ~5x as many linear adhesions formed on the few kPa substrate as the sub kPa substrate (fig. 3C vs 3A), whereas ~2x as many irregular adhesions formed on the sub kPa substrate as the few kPa substrate (fig. 3B vs 3D). This shows that E-cadherin adhesions can differentially sense sub kPa vs few kPa elasticity and organize differently in response. Notably, when we integrated the E-cadherin intensity in all the adhesions (both linear and irregular) across a similar number of cells on either substrate, we found that the total E-cadherin levels in all adhesions together were similar for the 0.3 kPa and 2.4 kPa substrate (fig. 3E). This suggests that about the same level of E-cadherin is distributed in adhesions in different ways in response to the sub kPa and few kPa elastic E-cadherin substrates.

**Figure 3.**
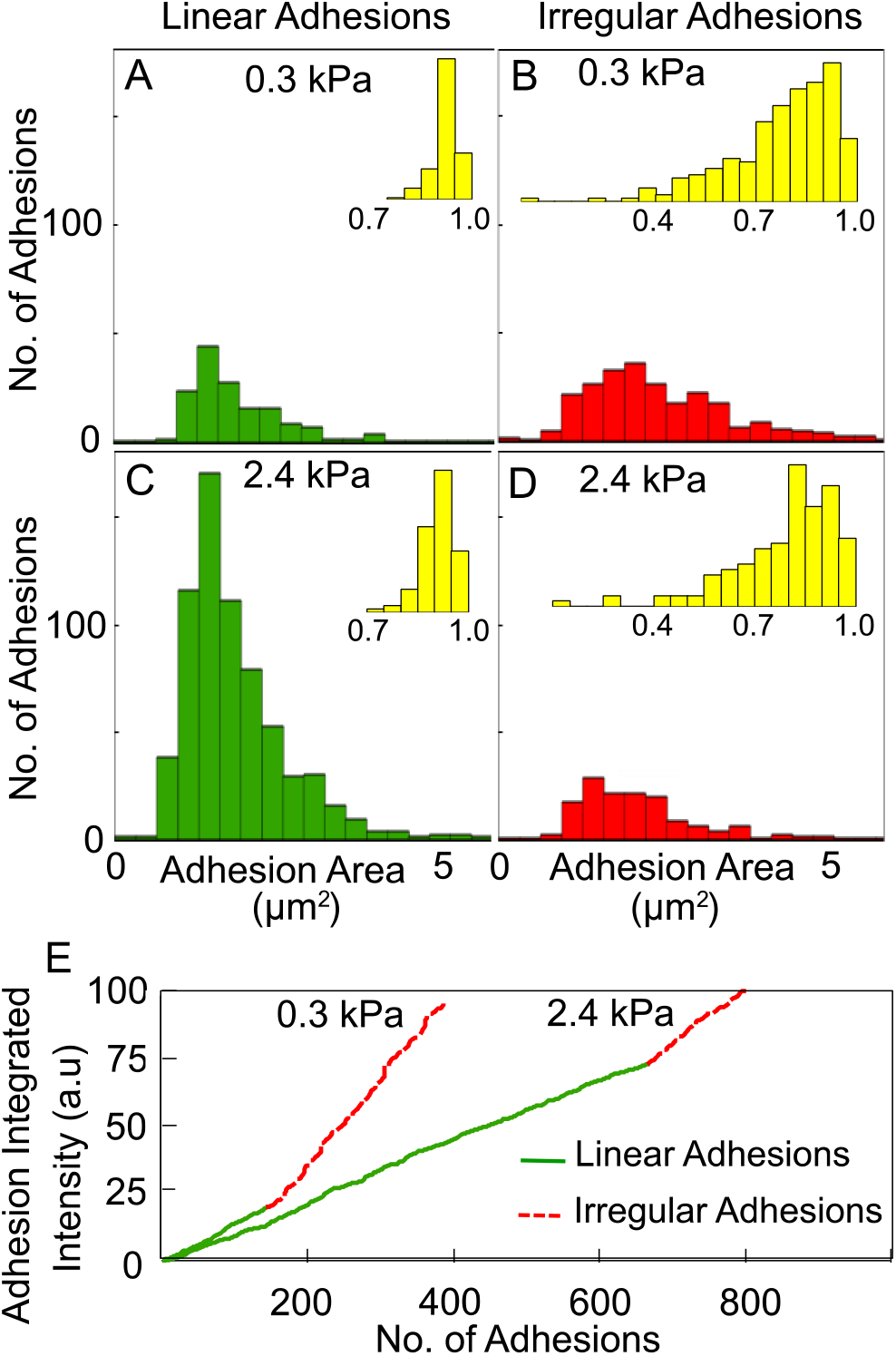
E-cadherin adhesion distribution depends on epithelial cell-like elasticity. (A-D) E-cadherin adhesion number versus area for linear adhesions (A, C) and irregular adhesions (B, D) as a function of E-cadherin substrate elasticity - 0.3 kPa (A, B) or 2.4 kPa (C, D). *Inset, histograms in yellow (A-D)*: E-cadherin adhesion number versus eccentricity for linear (A, C) and irregular (B, D) adhesions. (E) Sum of the integrated intensity of all linear (green, solid) and irregular (red, dashed) E-cadherin adhesions on 0.3 kPa and 2.4 kPa E-cadherin substrates. Data pooled from 3 independent experiments, with a total of 58 cells at 0.3 kPa and 59 cells at 2.4 kPa.

Since we observed that E-cadherin adhesions were affected by epithelial cell-like elasticity, we wanted to ascertain if this E-cadherin mechanosensing of elasticity led to distinct downstream events. We were particularly interested in nuclear recruitment of the transcriptional activator Yes-associated protein (YAP), which has been identified as a key player in mechanosensing pathways in various contexts [38]. On the one hand, E-cadherin adhesion has been shown to stimulate the Hippo pathway and suppress YAP nuclear recruitment [39–41]. On the other hand, tension across adherens junctions has been shown to inhibit the Hippo pathway and promote YAP nuclear recruitment [42, 43]. Given that human epithelial cancer cells are often softer than normal cells, we wondered if E-cadherin sensing of epithelial cell-like elasticity affects YAP nuclear recruitment. We assessed the ratio of YAP intensity in the nucleus (segmented by DAPI stain) to that in the cytoplasm (outer boundary segmented using actin stain). As shown in fig. 4, we found that the average nuclear recruitment of YAP on the 0.3 kPa and 2.4 kPa E-cadherin substrates were not statistically different (p = 0.30, Welch’s t-test). This likely reflects an interplay of the effects of E-cadherin adhesion and tension transmission, both of which are downstream of microenvironment elasticity in our E-cadherin substrate system. However, it is notable that there was considerable variation in the YAP intensity ratios for the sub kPa substrate such that some of the cells displayed relatively high YAP nuclear recruitment.

**Figure 4.**
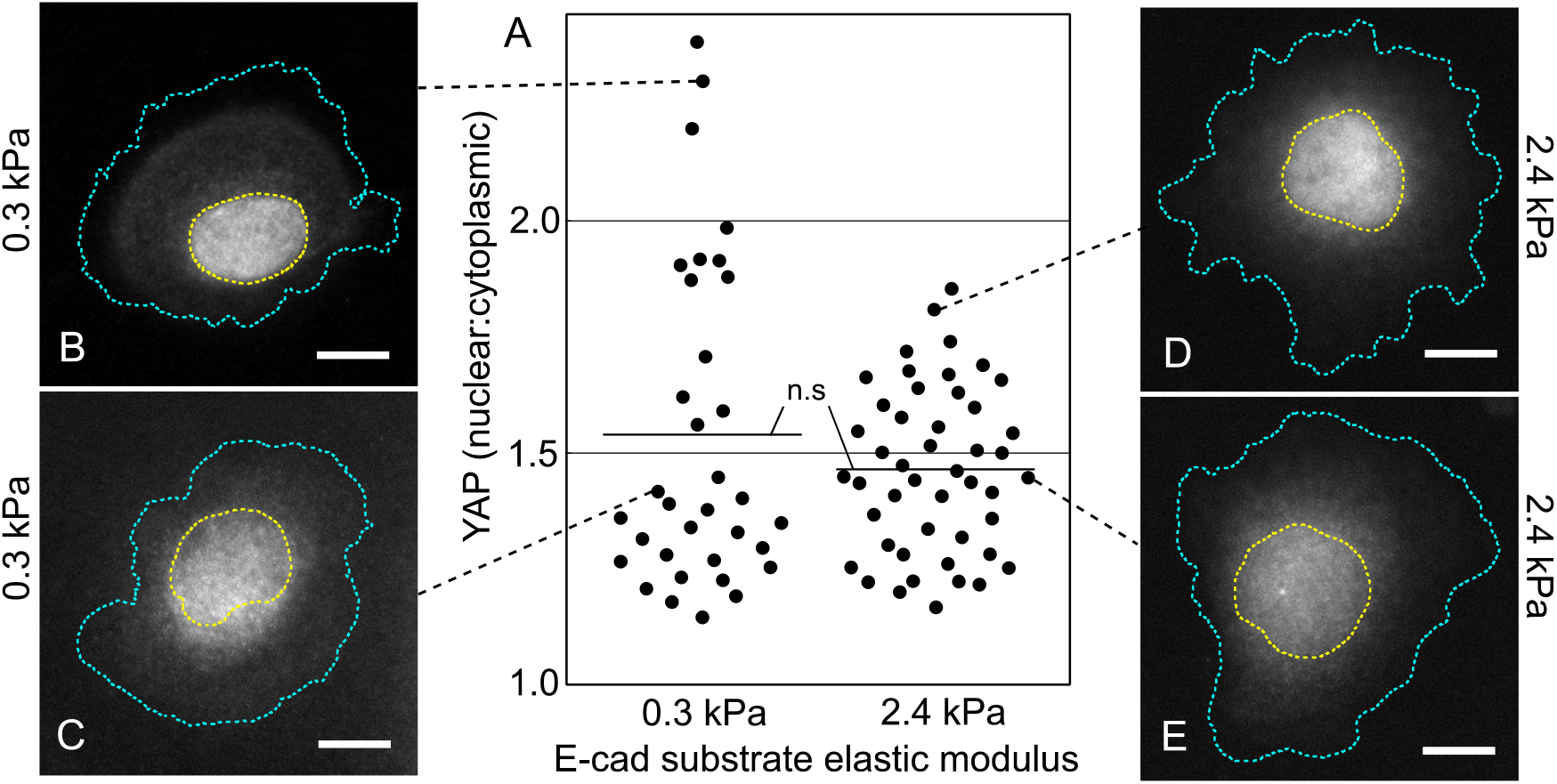
YAP nuclear recruitment on sub kPa and few kPa E-cadherin substrates. (A) Ratio of nuclear to cytoplasmic YAP intensity for 0.3 kPa and 2.4 kPa E-cadherin substrates. Short horizontal lines indicate the mean for each case. n.s. = not statistically significant. Data pooled from 3 independent experiments, with a total of 34 cells at 0.3 kPa and 47 cells at 2.4 kPa. (B-D) Representative immunofluorescence images of C2BBe cells stained for YAP, corresponding to the points indicated using the black, dotted lines. Yellow dotted curves enclose the nuclei and cyan dotted curves indicate the cell boundaries in (B-D). All scale bars are 10 μm.

In this report, we devised biomimetic E-cadherin soft substrates with epithelial-cell like elasticities and showed that E-cadherin adhesion distribution is different on sub kPa and few kPa substrates. It is worth noting that, since only E-cadherin-based adhesions link the cell to the substrate here, this involves E-cadherin-based sensing of substrate stiffness as well as the response of altered E-cadherin organization. It will be interesting to identify the extent to which various cellular factors (such as Rho GTPases and actin binding proteins) facilitate this specific effect of microenvironmental elasticity on E-cadherin adhesion organization. E-cadherin sensing of epithelial cell-like elasticity may have implications in various physiological contexts where epithelial cell stiffness changes, such as during wound healing [44]. In the context of cancer mechanobiology, this may be one factor that influences how cell stiffness changes (during cancer progression) impact cell-cell interactions. While we found that the average YAP nuclear recruitment is not different for sub kPa and few kPa E-cadherin substrate elasticities, how adhesion and force transmission specifically contribute to downstream events in this context remains to be teased out. E-cadherin adhesions serve a complex role – organizing cell-cell junctions, mechanically coupling neighboring cell cortices, maintaining cell polarity, and influencing a myriad of signaling pathways. We propose that E-cadherin sensing of epithelial cell-like stiffness may be an additional key role of these versatile adhesions.

## Acknowledgments

We thank Guijun Wang and Pooja Sharma for assistance with the rheology measurements. We thank Allen Ehrlicher for a useful discussion on soft silicone substrates. The web app PlotsOfData [45] was used to render the plot in figure 4. V.M. acknowledges support from the National Institutes of Health under award number 1R15GM116082.

## Author contributions

V.M. designed research; M.E. and S.C. performed research; M.E. and V.M. analyzed data; V.M. wrote the paper.

